# Recurrent inhibition, not presynaptic inhibition, contributes to the velocity-dependent control of motoneuron output during eccentric contractions

**DOI:** 10.64898/2026.07.20.739651

**Authors:** Julian Colard, Kazunori Nosaka, Christopher Latella, James O’Loughlin, Thomas Cattagni, Marc Jubeau

**Author notes:** CORRESPONDING AUTHOR: Marc Jubeau, PhD, Nantes University, Movement - Interactions - Performance, MIP, UR 4334, F-44000 Nantes, France, 25 bis Boulevard Guy Mollet - BP 72206, 44 322 Nantes cedex 3, France.

## Abstract

It is well documented that both motoneuron output and the effectiveness of activated Ia afferents to discharge soleus α-motoneurons decrease during eccentric (muscle lengthening) contractions. Evidence suggests that these modulations can be explained by recurrent inhibition and greater presynaptic inhibition of Ia afferents. However, the influence of angular velocity on the modulation of the effectiveness of activated Ia afferents to discharge α-motoneurons observed during eccentric contractions remains unclear. We investigated the influence of angular velocity on spinal mechanisms involved in the effectiveness of activated Ia afferents to discharge α-motoneurons during eccentric plantar flexor contractions using 16 healthy adults. We used both simple and conditioned Hoffmann reflex with different conditioning techniques to assess presynaptic inhibition, heteronymous Ia facilitation and heteronymous recurrent inhibition coupled with electromyography during eccentric contractions of the plantar flexors at three angular velocities. Our results showed that during eccentric contractions, the effectiveness of Ia afferents to discharge α-motoneurons was lower at 90°·s⁻¹ than 60°·s⁻¹ and 20°·s⁻¹ angular velocities. A similar velocity-dependent pattern was observed for heteronymous recurrent inhibition, decreasing at 90°·s⁻¹ when compared with 60°·s⁻¹ and 20°·s⁻¹. In contrast, presynaptic inhibition of Ia afferents was not different between the velocities. These demonstrate a differential influence of angular velocity on spinal recurrent inhibitory mechanisms during eccentric contractions and support distinct functional roles of recurrent and presynaptic inhibition in modulating α-motoneurons discharge with increasing movement velocity. The findings provide new insights into the velocity-dependent and mechanism-specific modulation of spinal inhibitory circuits during eccentric contractions.

**KEY POINTS:**

- During eccentric contractions in soleus muscle, the effectiveness of activated Ia afferents to discharge α-motoneurones decreases with increasing angular velocity, indicating a velocity-dependent modulation.
- Presynaptic inhibition of Ia afferents does not differ between angular velocities, suggesting that it does not contribute to the observed changes.
- Heteronymous recurrent inhibition from the quadriceps to the soleus increases with angular velocity, indicating that increasing movement velocity promotes a functional reorganization of intermuscular recurrent inhibition.
- These findings suggest a differential functional role of the two spinal inhibitory mechanisms, indicating that increasing angular velocity primarily influences recurrent postsynaptic inhibition rather than presynaptic inhibition.

## INTRODUCTION

Eccentric contractions represent muscle contractions in which active muscles are lengthened by greater external force than muscle force (Lindstedt *et al*., 2001). During maximal contractions, higher forces can be produced during eccentric than during isometric and concentric (shortening) contractions, while for a given absolute force, motoneuronal output and muscle activation are lower in eccentric contractions (Westing *et al*., 1991; Komi *et al*., 2000; Pasquet *et al*., 2000; Aagaard *et al*., 2000; Babault *et al*., 2003). This specific force-activation relationship suggests a distinct neural control strategy during eccentric contractions. This control is likely influenced by the conditions under which eccentric contractions are performed, particularly muscle length, as previously demonstrated (Colard *et al*., 2025), contraction intensity (Duclay *et al*., 2011), and potentially contraction velocity. However, the influence of angular velocity on neural control during eccentric contractions remains unclear despite its importance for understanding the regulation of this type of contraction during functional tasks such as locomotion.

Current evidence suggests that inhibitory mechanisms acting at the spinal level contribute strongly to the specific control of eccentric contractions (Gruber *et al*., 2009; Howatson *et al*., 2011; Duclay *et al*., 2014). At this level, the effectiveness of activated Ia afferents to discharge α-motoneurons (Ia-to-motoneuron transmission) is reduced compared to other contraction types in the soleus (SOL) muscle (Duclay & Martin, 2005). Among the mechanisms that may contribute to this reduction, recurrent inhibition mediated by Renshaw cell activity (Barrué-Belou *et al*., 2018, 2019; Papitsa *et al*., 2022) and primary afferent depolarization (PAD) reflecting presynaptic inhibition of Ia terminals (Colard *et al*., 2023), have been shown to be involved.

These findings suggest that spinal inhibitory circuits play a key role in modulating motoneuron output during eccentric contractions. However, recent work from our group (Colard *et al*., 2025) indicated that these modulations were not fixed but rather depended on the mechanical conditions under which the contraction was performed. In particular, muscle length was shown to influence neuromuscular responses during eccentric contractions, supporting the idea that neural control of eccentric actions is sensitive to mechanical context (Colard *et al*., 2025).

Contraction velocity represents another major mechanical factor likely to influence this control. Indeed, during muscle lengthening, both muscle length and stretch velocity strongly influence sensory feedback and reflex behaviour. In particular, changes in stretch velocity substantially alter afferent input and spinal excitability (Burke *et al*., 1978). Similarly, Pinniger *et al*. (2001) demonstrated that the amount of Ia-to-motoneuron transmission from SOL muscle, observed during passive muscle lengthening, decreased with greater angular velocity. In addition, the Ia-to-motoneuron transmission was decreased during eccentric but not concentric maximal and submaximal contractions at the angular velocity of 50 and/or 60°·s⁻¹ but not at the angular velocity of 20°·s⁻¹ (Romanò & Schieppati, 1987; Duclay *et al*., 2009). Although presynaptic mechanisms such as presynaptic inhibition have been suggested (Duclay *et al*., 2009), another study supports the hypothesis that sensory information arising from muscle spindle afferents plays a minor role in modulating the Ia-to-motoneuron transmission during eccentric contractions at different angular velocities (Valadão *et al*., 2018). Taken together, these suggest that neural control of eccentric contractions is velocity-dependent, but it remains unclear how specific inhibitory mechanisms are modulated across different angular velocities.

Most studies investigating spinal inhibitory mechanisms during eccentric contractions have been conducted at relatively low angular velocities, typically around 20°·s⁻¹. This limitation raises the question of whether these findings can be generalized to faster, more functional movement speeds. Such speeds are typically encountered during walking or running, where joint angular velocities of lower limb joints are substantially higher than 20°·s⁻¹. For example, during the mid-stance phase of walking, ankle dorsiflexion (i.e., eccentric contraction) occurs at angular velocities ranging from ∼44 to 93°·s⁻¹, at walking speeds of approximately 2.2 to 5.8 km·h⁻¹ (Mentiplay *et al*., 2018). Addressing this issue is therefore important for understanding how spinal inhibitory mechanisms operate under ecological conditions. While some studies (Kovaleski *et al*., 1995; Plautard *et al*., 2015) have primarily focused on biomechanical parameters related to contraction velocity, a comprehensive understanding of how spinal inhibitory mechanisms behave across a wider range of velocities remains lacking. This is particularly relevant given that most functional movements, such as locomotion, involve continuously changing rather than constant angular velocities (Mentiplay *et al*., 2018). Such conditions are notably encountered during gait initiation (i.e., the transition from standing to walking) or during gait acceleration, which require precise and rapid modulation of neural output.

Therefore, we aimed to examine the effects of ankle angular velocity on presynaptic and recurrent inhibition during eccentric contractions in SOL. We hypothesised that increased presynaptic inhibition and recurrent inhibition previously observed during eccentric contractions would be more pronounced at higher angular velocities (e.g., 90°·s⁻¹ compared with 60°·s⁻¹ and 20°·s⁻¹). By addressing this gap, we investigated how spinal inhibitory mechanisms would behave with variations in angular velocity during voluntary eccentric contractions.

## MATERIALS & METHODS

### Ethical approval

The study conformed with standards set by the Declaration of Helsinki (2013), except for registration in a database, and was approved by the Human Research Ethics Committee of the Edith Cowan University (2025-06296). All volunteers in the study gave their informed written consent before participation in the study.

### Participants

We used the G*Power software (version 3.1.9.6) to determine the minimum sample size required for assessing the primary expected outcomes. This calculation was based on a prior investigation (Colard et al., 2025) with a power (1 − β) of 0.95, an α of 0.05, and η^2^_p_ (partial η ^2^) values of 0.30 for presynaptic inhibition and 0.50 for spinal recurrent inhibition. The analysis suggested that a minimum of 8 participants would be necessary for presynaptic inhibition and spinal recurrent inhibition in our selected statistical analysis model (repeated measures ANOVA). However, to accommodate potential dropouts, we recruited 16 participants with no history of neurological illness or musculoskeletal injury to the ankle plantarflexors. Their average ± SD (range) age, height and body mass were 23.7 ± 4.1 (18–32) years, 172.6 ± 6.6 (160–186) cm, and 73.5 ± 20.1 (54–102) kg, respectively. Participants were instructed to refrain from strenuous exercise or resistance training during the week prior to and throughout the experimental period. In addition, they were asked to abstain from caffeine consumption on the day of each experimental session.

### Study design

All 16 participants took part in two separate experimental sessions. In each session, assessments of eccentric contraction at different angular velocities (i.e., 20°·s⁻¹, 60°·s⁻¹, and 90°·s⁻¹) were randomly assigned to avoid potential bias related to testing order. These velocities were selected based on methodological and physiological considerations, including the need to ensure reliable measurements and consistent stimulation triggering at a common joint ankle angle across conditions. They were also chosen to maximize the likelihood of detecting differences between velocities while remaining within a functionally relevant range, approximating those encountered during walking (Mentiplay *et al*., 2018).

In the first session (Experiment A), the Hoffmann reflex (H-reflex) and M-wave recruitment curves in the SOL muscle were assessed. In the second session (Experiment B), the effect of angular velocity on presynaptic and recurrent inhibition was investigated using the D1 method, heteronymous Ia facilitation (HF) and the heteronymous recurrent inhibition method. All experimental data were collected from participants’ right leg.

### Experimental set-up

#### Dynamometer

The participants were seated with the trunk inclined at 30°, the hip flexed at 85° (0° representing the anatomical position), and the knee joint at 0° of flexion (0° representing full extension). A Biodex ergometer enabled instantaneous recording of ankle angle and angular velocity during eccentric contractions (Biodex 3, Shirley, NY, USA). The output signal from the dynamometer was collected using a commercial acquisition system (CED, Micro1401-4, Cambridge Electronic Design; Cambridge, UK), displayed, and stored using Spike2 (Version 10; Cambridge Electronic Design; Cambridge, UK).

During eccentric contractions, the initial ankle angle varied depending on the angular velocity tested (see the percutaneous electrical stimulation section for more details). During the two experiments, each dynamometer movement cycle lasted 6.2 seconds and consisted of 1s of pre-activation, 0.6 seconds of active movement (dorsiflexion), 0.6 seconds to return to the initial position, and 4 seconds of rest before the next cycle (Table 1). Prior to each measurement, the ankle was maintained at rest for 4 seconds in the starting position. Participants first performed an isometric pre-activation of the plantar flexors for 1 second at 30% of their maximal electromyographic (EMG) activity. This step ensured consistent thixotropic effects across all test conditions, as described by (Proske *et al*., 1993).

**Table 1:** Experimental parameters for each angular velocity during eccentric contractions. °, degree; °·s⁻¹, degree per second; s, second; PF, plantarflexion; DF, dorsiflexion.

|  | Pre activation time | Initial angle (PF,+) | Final angle (DF, -) | Contraction time | Return time | Rest | Total time |
| --- | --- | --- | --- | --- | --- | --- | --- |
| **20°·s<sup>-1</sup>** | 1 s | 6° | -6° | 0.6 s | 0.6 s | 4 s | 6.2 s |
| **60°·s<sup>-1</sup>** | 1 s | 18° | -18° | 0.6 s | 0.6 s | 4 s | 6.2 s |
| **90°·s<sup>-1</sup>** | 1 s | 27° | -27° | 0.6 s | 0.6 s | 4 s | 6.2 s |

#### Electromyography

Surface EMG signals were recorded from the SOL, tibialis anterior (TA) and vastus lateralis (VL) muscles using pairs of self-adhesive surface electrodes in a bipolar configuration (Meditrace 100; Covidien, Mansfield, MA, USA). SOL electrodes were placed 2 cm below the muscle–tendon junction of the gastrocnemii. For the TA, electrodes were placed on the muscle belly, parallel to the longitudinal axis of the muscle, at one-third of the distance between the head of the fibula and the tip of the medial malleolus. VL electrodes were placed at a position two-thirds along the line from the anterior superior iliac spine to the lateral side of the patella.

The skin was abraded and cleaned with alcohol. EMG signals were amplified (×1000), filtered (10 Hz–1 kHz; CED 1902 amplifier, Cambridge Electronic Designs, Cambridge, UK), digitized at 2 kHz (CED Micro1401-4, Cambridge Electronic Designs, Cambridge, UK) and stored for offline analysis using Spike2 (Version 9).

#### Percutaneous electrical nerve stimulation

The electrophysiological responses (i.e., H-reflex and M-wave) of the SOL were evoked by percutaneous electrical stimulation of the posterior tibial nerve using a single rectangular pulse (1 ms) and high voltage (400 V), delivered by a stimulator (Digitimer, model DS7R, Constant Current Stimulator; Hertfordshire, UK). A self-adhesive electrode (1 cm diameter, Ag–AgCl) was used as the cathode and was placed over the popliteal fossa. The anode electrode was placed on the anterior surface of the knee, below the patella. Once the optimal position was determined, the cathode electrode was firmly secured with a strap and tape.

To assess presynaptic inhibition through primary afferent depolarization modulation (D1 method), the fibular nerve was stimulated using a second DS7A current stimulator (Digitimer; Hertfordshire, UK), with the cathode electrode placed close to the head of the fibula and the anode electrode near the medial part of the tibial head. For heteronymous Ia facilitation and recurrent inhibition, the femoral nerve was stimulated with the cathode positioned over the nerve in the femoral triangle and the anode placed over the greater trochanter. The stimulation site was first determined by obtaining an M-wave in the VL associated with an upward movement of the patella. The motor thresholds for fibular and femoral nerve stimulation were defined as the lowest intensity that evoked at least three M-waves out of five stimulations.

To ensure a rigorous comparison of the effect of angular velocity on H-reflex modulation, it was crucial to maintain a constant delay between the onset of eccentric movement and the stimulation. As highlighted by Pinniger *et al*. (2001) and Robertson *et al*. (2013), the time interval between movement initiation and stimulation influenced H-reflex modulation more strongly than angular velocity itself. To isolate the effect of velocity, stimulation was applied at a fixed joint position (0°) with a consistent delay of 333 ms after movement onset (Budini & Tilp, 2016). Accordingly, the starting and finishing angles were adjusted for each angular velocity to maintain this constant delay. The movement ranged from 27° (plantar flexion) to −27° (dorsiflexion), from 18° to −18° and from 6° to −6°, at 90°·s⁻¹, 60°·s⁻¹, and 20°·s⁻¹, respectively (Table 1).

### Experimental protocol

#### Experiment A

First, three maximal voluntary eccentric contractions at 60°·s⁻¹ were performed to determine the maximal SOL EMG root mean square (RMS) level, calculated using a 500-ms moving window. The maximum SOL EMG RMS activity recorded during eccentric MVCs was used to establish the target activation level (30% of maximal SOL EMG RMS) for participants during the study. Participants were asked to contract their plantarflexors so that they moved and maintained their SOL EMG RMS biofeedback on the target corresponding to 30% of maximal EMG RMS. The biofeedback was provided by a specific digital computing channel. This channel instantaneously computed the RMS level of the amplified EMG signal with an integration time of 500 ms. For each submaximal contraction, participants were asked to reach the target and maintain it during the entire duration of the contraction. This method ensured that muscle activation levels remained consistent across all test conditions (Duclay *et al*., 2014; Colard *et al*., 2023).

Second, the maximal H-reflex (H_max_) and maximal M-wave (M_max_) were measured at each angular velocity (20°·s⁻¹, 60°·s⁻¹ and 90°·s⁻¹) during eccentric contractions. SOL H-reflex and M-wave recruitment curves were determined during plantarflexion at 30% of maximal EMG RMS, resulting in a total of three recruitment curves. The stimulation intensity was gradually increased in 2-mA increments from the H-reflex threshold to the intensity at which the maximal H-reflex was recorded for SOL. Then, the intensity was increased with a 5-mA increment until the intensity for which no further increase in the amplitude of the M-wave (i.e., plateau) was observed. Finally, supramaximal stimulations at 120% of this latter stimulation intensity were delivered to ensure the recording of SOL M_max_. Five electrical stimulations were delivered at each stimulation intensity, as recommended in literature (Hopkins *et al*., 2000; Theodosiadou *et al*., 2023).

#### Experiment B

To investigate presynaptic inhibition, we used the D1 method, as described by Mizuno *et al*. (1971). This technique involved eliciting an electrical volley in the nerve of the antagonist muscle, thereby activating Ia afferent fibres, which in turn induce presynaptic inhibition of the Ia terminals projecting to the α-motoneuron pool, resulting in a depression of the SOL H-reflex (Pierrot-Deseilligny & Burke, 2005). Recent paradigms have suggested that presynaptic inhibition is triggered by GABAergic interneurons located near dorsal nodes of Ranvier (Hari *et al*., 2022). While PAD can facilitate Ia afferent conduction through the generation of PAD-evoked spikes, excessive PAD may ultimately produce a presynaptic inhibitory effect, contributing to H-reflex depression (Metz *et al*., 2023).

SOL H_test_ was conditioned with a stimulus applied to the fibular nerve, which activated GABAergic interneurons responsible for primary afferent depolarization of Ia afferents from the SOL muscle. To evoke the SOL conditioned H-reflex (H_D1_), a single 1-ms stimulus was applied to the fibular nerve, with an intensity equivalent to 1.2 times the tibialis anterior motor threshold (Howells et al., 2020). The interval between the stimulation and the SOL H_test_ was 21 ms (Aymard *et al*., 2000; Magalhães *et al*., 2015). 20 H_D1_ and 20 H_test_ were randomly evoked during eccentric contractions at each angular velocity (20°·s⁻¹, 60°·s⁻¹ and 90°·s⁻¹)

To estimate the level of heteronymous Ia facilitation, the method of Hultborn *et al*. (1987) was used, which evaluates the monosynaptic facilitatory effect of femoral nerve stimulation on the amplitude of the SOL H-reflex. To minimize contamination from polysynaptic inputs (e.g., Ib projections), femoral nerve stimulation followed tibial nerve stimulation due to the shorter neural pathway of the heteronymous Ia afferent pathway compared with the homonymous pathway. By considering the mean duration of a synaptic event (0.5 to 1 ms) we aimed to minimize the influence of polysynaptic excitatory inputs (Baudry & Enoka, 2009; Souron *et al*., 2019). For each participant, the onset of H-reflex facilitation in response to femoral nerve stimulation was determined by adjusting the delay between the test (tibial nerve) and conditioning (femoral nerve) stimuli in 0.5-ms increments, ranging from 9 ms (test stimulus preceding the conditioning stimulus by 9 ms) to 1 ms (test stimulus preceding the conditioning stimulus by 1 ms). The conditioning stimulation intensity was set at 1.3 times the vastus lateralis motor threshold, and an average of at least five trials was conducted for each delay. The criterion used to identify facilitation onset was an increase in conditioned H-reflex (H_Fac_) amplitude of more than 10% over non-conditioned H-reflexes (Johannsson *et al*., 2015). A total of 20 H_Fac_ and 20 H_test_ were randomly evoked in each condition.

To assess recurrent inhibition, we employed the heteronymous recurrent inhibition method. The heteronymous recurrent inhibition paradigm was used to investigate the functional organization of recurrent spinal networks across muscles rather than local recurrent feedback within a single motoneuron pool (Barbeau *et al*., 2000). This approach also reduces the influence of methodological confounders inherent to homonymous paradigms, including post-activation depression and other stimulation-related effects (Mazzocchio & Rossi, 1989; Rossi & Mazzocchio, 1991). Test stimuli were applied to the posterior tibial nerve, and conditioning stimuli were applied to the femoral nerve to assess the level of recurrent inhibition produced in SOL motoneurons by activation of the recurrent collaterals of the quadriceps motor axons (H_RI_). The intensity of the test stimuli was adjusted to produce an H-reflex of approximately 25% of M_max_ (H_test_) in the SOL EMG (Sangari *et al*., 2022), while the intensity of the conditioning stimuli was set at the threshold required to produce M_max_ in the vastus lateralis EMG. The interval between femoral and posterior tibial nerve stimuli was set at 15 ms, which has been reported to be optimal for recurrent inhibition in SOL motoneurons (Barbeau *et al*., 2000). The size of the conditioning M_max_ was monitored throughout the experiment to ensure the stability of the conditioning stimuli. A total of 20 H_RI_ and 20 H_test_ were randomly evoked in each condition.

### Data analyses

#### Evoked potentials

For each angular velocity, the mean peak-to-peak amplitude of SOL H_max_, M_max_, H_D1_, H_FAC_, H_RI_ and H_test_ were calculated over all trials. All ratios of evoked potentials were calculated during eccentric contractions at each angular velocity (20, 60 and 90°·s⁻¹).

To estimate the Ia-to-motoneuron transmission, we calculated H_max_/M_max_. This ratio is commonly used as an indicator of the maximal proportion of α-motoneurons recruited by the Ia afferent pathway. The M-wave elicited concomitantly with the H_max_ (M_at_H_max_), a small fraction of the M_max_ (M_at_H_max_/M_max_), was measured and analysed to verify the stability of the stimulus intensity in each corresponding condition (Schieppati, 1987). To assess presynaptic inhibition of Ia afferents, we expressed H_D1_ as a percentage of mean H_test_. A higher percentage value indicates lower presynaptic inhibition of Ia afferents. The H_test_/M_max_ ratio was monitored to ensure that the amplitude of the non-conditioned H-reflex remained consistent across experimental conditions.

To assess heteronymous Ia facilitation, we calculated H_FAC_ as a percentage of H_test_. A higher H_FAC_/H_test_ reflects greater facilitation (Baudry & Enoka, 2009). To assess recurrent inhibition, we expressed H_RI_ as a percentage of H_test_. In this case, a higher H_RI_/H_test_ ratio corresponds to less recurrent inhibition (Sangari *et al*., 2022).

#### EMG activity during voluntary contractions

RMS of the SOL EMG signal over a 500-ms period prior to stimulation was normalized to the corresponding RMS of M_max_ (EMG RMS/EMG RMS_Mmax_) during all conditions. The normalization procedure was used to ascertain whether SOL EMG activities remained constant across all experimental conditions. In addition, if the EMG RMS value of the SOL exceeded two standard deviations from the reference mean (considered to represent 30% of maximal EMG level), the evoked response following this deviation was excluded.

Similarly, RMS of the TA EMG signal was assessed over the same time period preceding the stimulation to check that the co-activation was constant. To assess the co-activation index between SOL and TA, we adapted the formula of (Banks *et al*., 2017):

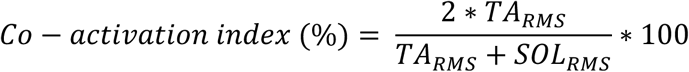

This index represents a ratio of agonist (SOL RMS)/antagonist (TA RMS) overlap to total muscle activation (SOL RMS + TA RMS). A co-activation index value of 100% represents total co-activation, while a value of 0% represents pure agonist (SOL) activation.

### Statistical analyses

All descriptive statistics presented in the text, tables and figures are provided as mean values ± standard deviation. The significance level for all analyses was set at P < 0.05. The normality of the data and homogeneity of variances were verified and validated using the Shapiro–Wilks W test and Levene test, respectively. Separate one-way repeated measures ANOVAs were used to compare SOL M_at_H_max_/M_max_, H_max_/M_max_, H_test_/M_max_, H_D1_/M_max_, H_D1_/H_test_, H_FAC_/M_max_, H_Fac_/H_test_, H_RI_/M_max_ and H_RI_/H_test_ during eccentric submaximal contractions at different angular velocity (20, 60 and 90°·s⁻¹). Two-factor [angular velocity (20°·s⁻¹ vs. 60°·s⁻¹ vs. 90°·s⁻¹) × experiment (experiment A vs. experiment B)] repeated measures ANOVAs on angular velocity and experiment were used to compare M_max_, SOL and TA EMG RMS/EMG RMS_Mmax_ and co-activation index. Whenever a significant main effect or interaction was detected, Tukey tests were performed for post hoc analysis. Statistical analyses were conducted using JASP (Version 0.17.2.1; Amsterdam, The Netherlands), while data visualization and figure illustration were performed using GraphPad Prism (Version 10.2.1; San Diego, CA, USA) and BioRender.

## RESULTS

### EMG activity

No significant effects were found for SOL EMG RMS/EMG RMS_Mmax_ (all P values > 0.48). No effect between conditions was found for the co-activation index (P values > 0.224). The mean co-activation index value was 10.3 ± 3.2% during all eccentric contractions.

### Ia-to-motoneuron transmission

Figure 1A shows representative raw traces of the SOL H-reflex and M-wave recorded during eccentric contractions performed at the three angular velocities. A significant main effect of contraction velocity was found for SOL H_max_/M_max_ (F_(2, 30)_ = 40.388; P < 0.001; ηp² = 0.729; Fig. 1B). During eccentric contractions at 90°·s⁻¹, SOL H_max_/M_max_ was reduced by 33.4% and 22.6% compared with 20°·s⁻¹ and 60°·s⁻¹, respectively (both P < 0.001). At 60°·s⁻¹, SOL H_max_/M_max_ was decreased by 14.2% relative to 20°·s⁻¹ (P < 0.001). No significant effect of contraction velocity was found for either SOL or VL M_max_ (both P = 0.112).

**Figure 1:**
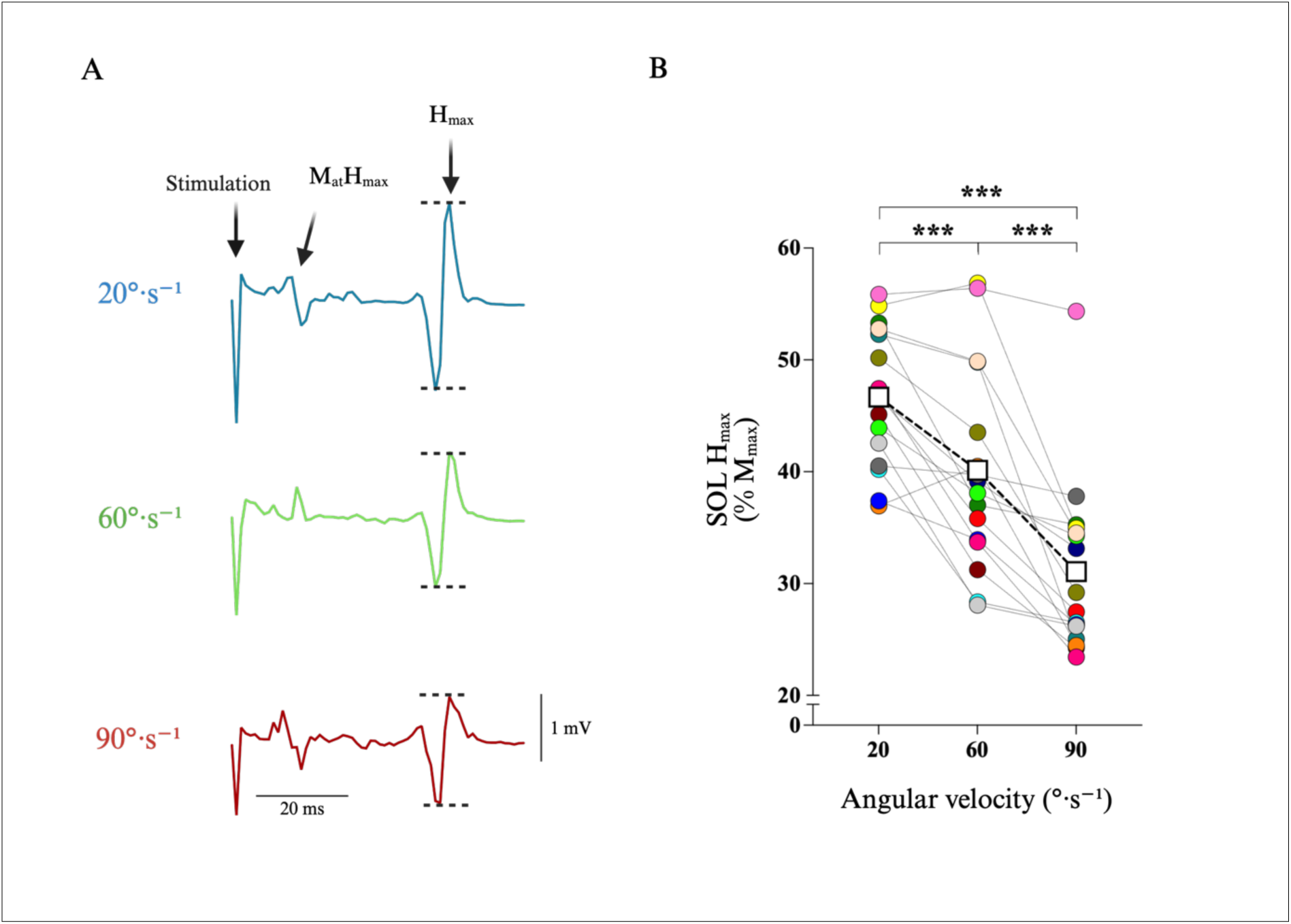
Changes in motor evoked potentials in soleus during eccentric contractions at 20, 60 and 90°·s⁻¹. A) Representative traces showing the maximal SOL H-reflex evoked by posterior tibial nerve stimulation during eccentric contractions at 20°·s⁻¹ (blue trace), 60°·s⁻¹ (green trace) and 90°·s⁻¹ (red trace). Black dashed lines indicate the maximal H-reflex amplitudes at each angular velocity during eccentric contractions. B) Absolute and individual data (n = 16), expressed as mean ± standard deviation. Changes in the maximal SOL H-reflex normalized to the corresponding maximal M-wave during eccentric contractions at 20, 60 and 90°·s⁻¹. SOL, soleus; H_max_, maximal SOL H-reflex; M_max_, maximal M-wave. *** Significantly different at P < 0.001.

During our experiment, we did not find an effect of angular velocity for SOL M_at_H_max_/M_max_. Our findings indicate that the amplitude of SOL M_at_H_max_/M_max_ remained consistent, suggesting that an equivalent proportion of α-motoneurons were activated across all conditions (Table 2). Thus, H_max_/M_max_ modulation cannot be attributed to recording conditions but rather to neural mechanisms (Romanò & Schieppati, 1987; Duclay & Martin, 2005).

**Table 2:** Effect of angular velocity on soleus and vastus lateralis evoked potentials. Absolute data (n = 16) are expressed as mean ± standard deviation. SOL, soleus; VL, vastus lateralis; M_max_, maximal M-wave; M_at_H_max_, M wave corresponding to maximal SOL H-reflex; H_test_, test SOL H-reflex; H_D1_, conditioning H-reflex amplitude for presynaptic inhibition; H_Fac_, conditioning H-reflex amplitude for heteronymous facilitation; H_RI_, conditioning H-reflex amplitude for heteronymous recurrent inhibition.

| | $20^{\circ}\cdot\text{s}^{-1}$ | $60^{\circ}\cdot\text{s}^{-1}$ | $90^{\circ}\cdot\text{s}^{-1}$ |
| --- | --- | --- | --- |
| **SOL $M_{\max}$ (mV)** | $3.810 \pm 0.956$ | $3.639 \pm 0.950$ | $3.743 \pm 0.827$ |
| **VL $M_{\max}$ (mV)** | $7.768 \pm 0.809$ | $7.355 \pm 0.798$ | $7.792 \pm 0.782$ |
| **$M_{\text{at}H_{\max}}/M_{\max}$ (%)** | $9.776 \pm 3.761$ | $8.900 \pm 3.331$ | $9.225 \pm 2.917$ |
| **$H_{\text{test}}/M_{\max}$ (%)** | $25.797 \pm 2.789$ | $26.244 \pm 2.240$ | $26.436 \pm 3.116$ |
| **$H_{D1}/M_{\max}$ (%)** | $15.630 \pm 4.860$ | $16.015 \pm 4.458$ | $15.085 \pm 4.256$ |
| **$H_{\text{Fac}}/M_{\max}$ (%)** | $34.149 \pm 4.971$ | $35.370 \pm 3.350$ | $34.988 \pm 4.473$ |
| **$H_{RI}/M_{\max}$ (%)** | $16.096 \pm 5.040$ | $13.458 \pm 3.571$ | $9.988 \pm 3.571$ |

### Presynaptic inhibition of Ia afferents (D1 method)

Figure 2A illustrates the experimental design used to estimate presynaptic inhibition from Ia afferents. Figure 2B shows representative traces of the conditioned (H_D1_) and test (H_test_) H-reflexes recorded from the SOL muscle during eccentric contractions performed at the three angular velocities. SOL H_D1_/H_test_ was used as an index of presynaptic inhibition of Ia afferents. No significant effect of contraction velocity was observed on SOL H_D1_/H_test_ (P = 0.167; Fig. 2C). Presynaptic inhibition was similar, with a H_D1_/H_test_ ratio of 58.2 ± 2.6%. Similarly, SOL H_D1_/M_max_ and SOL H_test_/M_max_ remained stable across all tested angular velocities (all P > 0.476).

**Figure 2:**
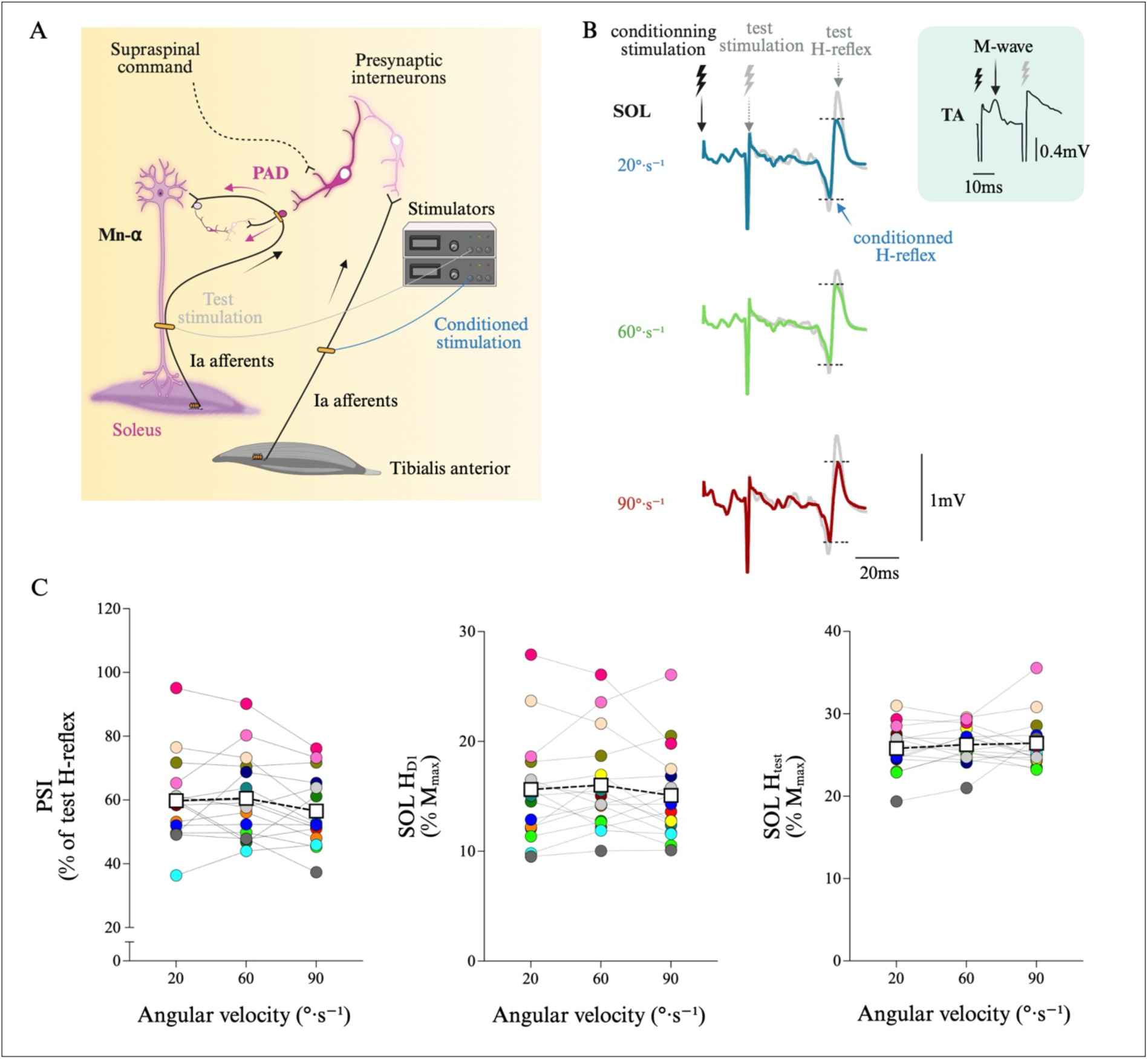
Changes in primary afferent depolarization during eccentric contraction at 20, 60 and 90°·s⁻¹. A) Schematic illustration of the experimental design used to assess presynaptic inhibition of Ia afferents using the D1 method. Conditioning stimulation (blue) of the antagonist nerve activates tibialis anterior Ia afferents, which induce primary afferent depolarization (PAD; purple) via activation of specific receptors, leading to presynaptic inhibition of SOL Ia terminals and a reduction in SOL H-reflex amplitude. B) Representative traces showing the test H-reflex (H_test_) and the conditioned SOL H-reflex (H_D1_) evoked by fibular nerve stimulation preceding posterior tibial nerve stimulation, with a conditioning test interval of 21 ms, during eccentric contractions at 20, 60 and 90°·s⁻¹. The stimulation intensity used to evoke the test H-reflex was normalized across all conditions. C) Absolute and individual data (n = 16) are expressed as mean ± standard deviation. The conditioned H-reflex and test H-reflex are expressed as a ratio during eccentric contractions at 20, 60, and 90°·s⁻¹. PSI, presynaptic inhibition of Ia afferents; SOL, soleus; TA, tibialis anterior; H_D1_, conditioned SOL H-reflex; H_test_, test SOL H-reflex; M-wave, maximal M-wave; PAD, primary afferent depolarization.

### Heteronymous Ia facilitation

Figure 3A illustrates the experimental design used to estimate heteronymous Ia facilitation. Figure 3B shows representative traces of the conditioned (H_Fac_) and test (H_test_) H-reflexes recorded from the SOL muscle during eccentric contractions performed at the three angular velocities tested. SOL H_Fac_/H_test_ was used as an index of Ia afferent facilitation. No significant effect of contraction velocity was observed on SOL H_Fac_/H_test_ (P = 0.482; Fig. 3C). Heteronymous Ia facilitation was similar, with a H_FAC_/H_test_ ratio of 131.4 ± 3.3%. Similarly, SOL H_Fac_/M_max_ remained stable across all tested angular velocities (P = 0.536).

**Figure 3:**
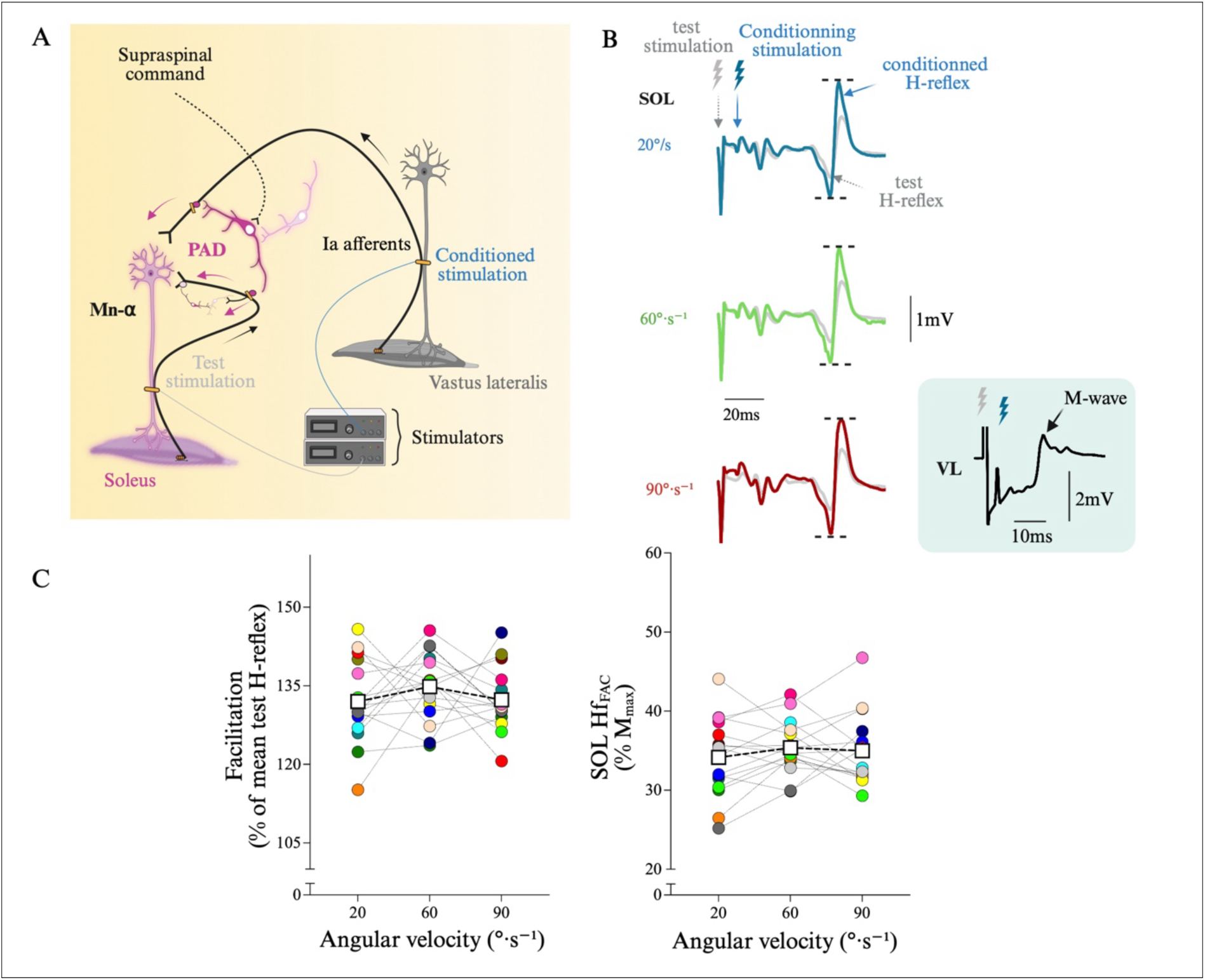
Changes in heteronymous Ia facilitation during eccentric contraction at 20, 60 and 90°·s⁻¹. A) Schematic illustration of the experimental design used to assess heteronymous Ia facilitation. Conditioning stimulation (blue) of the femoral nerve activates quadriceps Ia afferents, which facilitate SOL motoneuron activity via excitatory projections, leading to an increased SOL H-reflex amplitude (facilitation). These projections are also modulated by interneurons specifically mediating presynaptic inhibition. B) Representative traces showing the test H-reflex (H_test_) and the conditioned SOL H-reflex (H_FAC_) evoked by posterior tibial nerve stimulation preceding femoral nerve stimulation at a conditioning test interval ranging from 9 to 1 ms (0.5 ms step) during eccentric contractions at 20, 60 and 90 °·s⁻¹. The electrical intensity to evoke H_test_ was normalized during all conditions. C) Absolute and individual data (n =16) are expressed as mean ± standard deviation. The conditioned H-reflex and test H-reflex are expressed as a ratio during eccentric contractions at 20, 60, and 90°·s⁻¹. SOL, soleus; H_Fac_, conditioned SOL H-reflex; H_test_, test SOL H-reflex.

### Heteronymous recurrent inhibition

Figure 4A illustrates the experimental design used to estimate heteronymous recurrent inhibition in SOL muscle. Figure 4B, shows representative traces of the conditioned (H_RI_) and test (H_test_) H-reflexes recorded from the SOL muscle during eccentric contractions performed at the three angular velocities. A significant main effect of contraction velocity was found for SOL H_RI_/H_test_ (F_(2, 30)_ = 20.697; P < 0.001; ηp² = 0.664; Fig. 4C). During eccentric contractions at 90°·s⁻¹, SOL H_RI_/H_test_ was reduced by 39.1% and 26.8% compared with 20°·s⁻¹ and 60°·s⁻¹, respectively (both P < 0.001). At 60°·s⁻¹, SOL H_RI_/H_test_ was decreased by 16.8% relative to 20°·s⁻¹ (P = 0.011). Regarding SOL H_RI_/M_max_, we also found a significant effect of angular velocity (F_(2, 30)_ = 18.604; P < 0.001; ηp² = 0.554). During eccentric contractions at 90°·s⁻¹, SOL H_RI_/M_max_ was reduced by 37.9% and 25.8% compared with 20°·s⁻¹ and 60°·s⁻¹, respectively (P < 0.001 and P = 0.002). At 60°·s⁻¹, SOL H_max_/M_max_ was decreased by 16.4% relative to 20°·s⁻¹ (P = 0.027).

**Figure 4:**
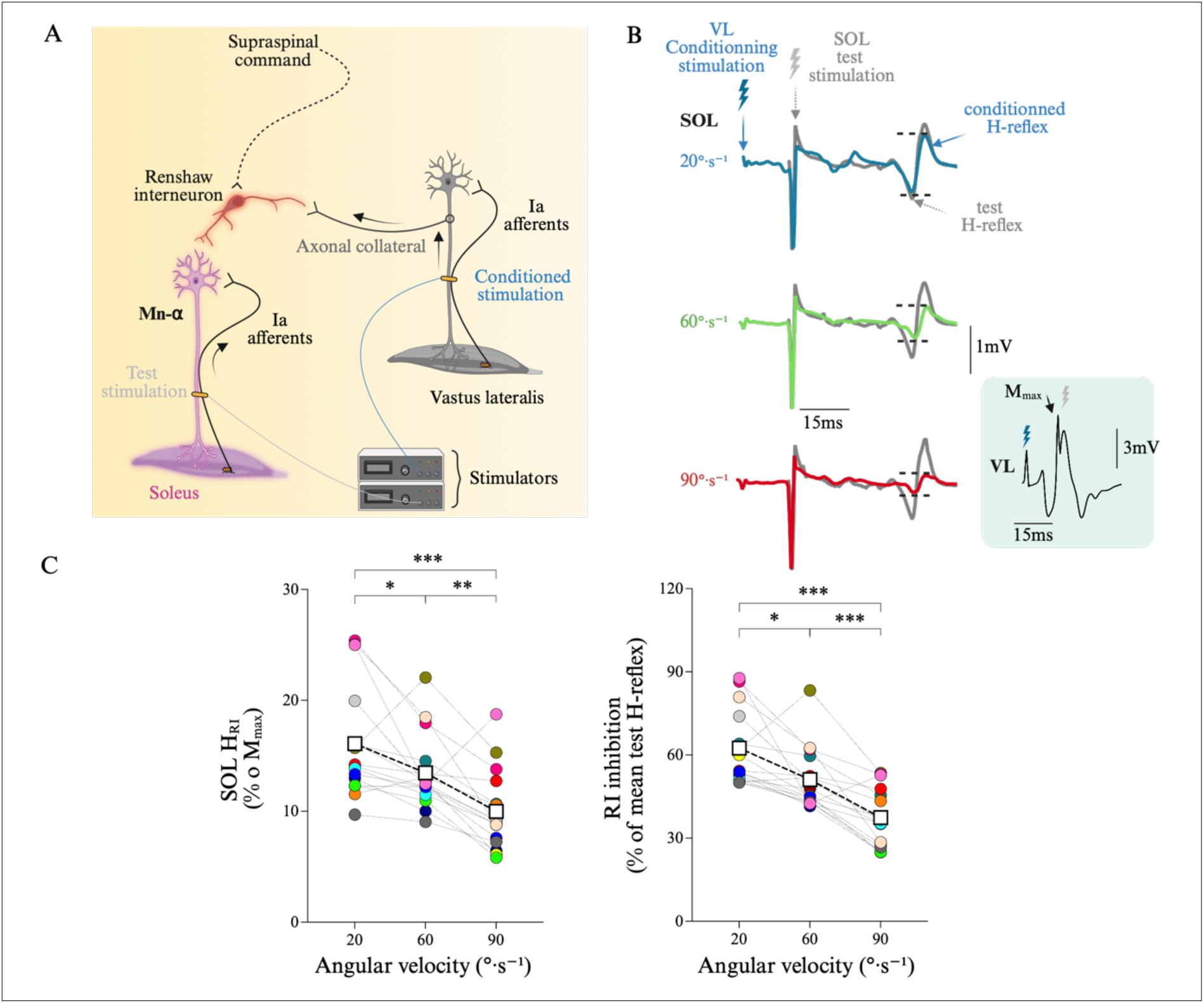
Changes in heteronymous recurrent inhibition during eccentric contraction at 20, 60 and 90°·s⁻¹. A) Schematic illustration of the experimental design used to assess heteronymous recurrent inhibition. Conditioning stimulation (blue) of the femoral nerve activates quadriceps motor axons, whose antidromic activity recruits Renshaw cells (red) via recurrent axon collaterals. These interneurons project onto motoneurons of heteronymous muscles, including SOL, resulting in a reduced SOL H-reflex amplitude. B) Representative traces showing the test H-reflex (H_test_) and the conditioned SOL H-reflex (H_RI_) evoked by femoral nerve stimulation preceding posterior tibial nerve stimulation at a conditioning test interval of 15 ms during eccentric contractions at 20, 60 and 90 °·s⁻¹. C) Absolute and individual data (n =16) are expressed as mean ± standard deviation. The conditioned H-reflex and test H-reflex are expressed as a ratio during eccentric contractions at 20, 60 and 90°·s⁻¹. B). The electrical intensity to evoke H_test_ was normalized during all conditions. SOL, soleus; VL, vastus lateralis; H_RI_, conditioned SOL H-reflex; H_test_, test SOL H-reflex. *, significantly different at P < 0.05; **, significantly different at P < 0.01; ***, significantly different at P < 0.001.

## DISCUSSION

The aim of this study was to investigate the effects of ankle angular velocity on pre- and post-synaptic inhibitory mechanisms during eccentric contractions. Our results clearly demonstrated a velocity-dependent modulation of Ia afferent-to-motoneuron transmission during eccentric contractions, characterized by a progressive reduction as angular velocity increases (Figure 1). The results also identified spinal inhibitory mechanisms contributing to this velocity-dependent modulation. While presynaptic inhibition of Ia afferents remained unchanged across the eccentric contraction velocities (Figure 2 and 3), heteronymous recurrent inhibition was enhanced as angular velocity increased (Figure 4). These findings indicate that the progressive reduction in Ia afferent-to-motoneuron transmission observed with increasing eccentric contraction velocity is primarily related to greater recurrent inhibition and not to changes in presynaptic inhibition. Overall, our findings demonstrate that spinal inhibitory mechanisms are differentially influenced by increasing angular velocity, highlighting the role of recurrent inhibition in the functional organization of intermuscular spinal connectivity.

### Velocity-dependent reduction of Ia–to-motoneuron transmission during eccentric contractions

The present study examined the influence of angular velocity on the Ia-to-motoneuron transmission and underlying mechanisms during eccentric muscle contractions. Our results indicate that the typical reduction in the Ia-to-motoneuron transmission observed during eccentric contractions at 20°·s⁻¹ compared to isometric and concentric contraction types was further accentuated at 60°·s⁻¹ and 90°·s⁻¹ (Figure 1). Although several previous studies have reported this effect during maximal contractions performed at angular velocities between 20°·s⁻¹ and 60°·s⁻¹ (Romanò & Schieppati, 1987; Duclay *et al*., 2009), our findings demonstrate that this phenomenon also occurs during submaximal contractions and becomes more pronounced at higher angular velocities. Previous studies have suggested that the velocity-dependent decrease in Ia-to-motoneuron transmission amplitude could be partly explained by increased presynaptic inhibition of Ia afferents arising from peripheral mechanisms, possibly associated with greater muscle spindle discharge at higher stretch velocities (Duclay *et al*., 2009). However, more recent evidence challenges this interpretation. Valadão *et al*. (2018) demonstrated that muscle spindle-related mechanical variables, such as fascicle length and pennation angle, remained similar across eccentric contractions performed at different angular velocities. In addition, higher fascicle velocity did not produce proportionally larger H-reflex responses, as would be expected if spindle-derived Ia input were the primary determinant (Valadão *et al*., 2018).

Functionally, the greater reduction in the H_max_/M_max_ ratio at higher angular velocities during eccentric contractions (Figure 1) may reflect a task-dependent regulation of Ia-to-motoneuron transmission aimed at limiting reflex excitability when muscles are actively lengthened at faster velocities. Such modulation likely contributes to preventing excessive reflex-driven activation and stiffness, thereby promoting smoother force control and mechanical stability during rapid lengthening actions (Duclay *et al*., 2009). This inhibitory adjustment may also favor greater reliance on supraspinal commands over spinal reflex pathways, thereby enhancing movement accuracy and adaptability during dynamic eccentric tasks. These findings do not support a predominantly peripheral, spindle-mediated explanation. Rather, they suggest that the modulation observed at higher angular velocities likely reflects supraspinal influences on presynaptic inhibition at the spinal level, consistent with previous observations during slower eccentric contractions (Colard *et al*., 2023).

### Presynaptic inhibition on velocity-dependent changes

Our results show that presynaptic inhibition of Ia afferents during eccentric contractions remains unchanged across angular velocities, as indicated by a consistent H_D1_/H_test_ ratio (Figure 2). This indicates that peripheral modulations of PAD, acting through GABAergic interneurons and reducing reflex activity, is unlikely to be involved (Metz *et al*., 2023). Moreover, although supraspinal commands are known to influence these same interneuron pathways, this hypothesis is not supported by the D1 conditioning method in the present study. An alternative explanation is that this inhibitory pathway may have already been operating close to its maximal functional level under our experimental conditions. This ceiling effect may involve a near-maximal engagement of GABA_A_-mediated nodal PAD and GABA_B_-mediated terminal presynaptic inhibition along Ia afferents, thereby constraining the dynamic range of further presynaptic modulation. If so, additional supraspinal input or peripheral modulation would not necessarily produce further measurable changes. This ceiling effect would suggest a saturation of the underlying neural mechanism, thereby masking potential velocity-dependent or task-dependent adjustments in presynaptic control.

The complementary results between the D1 method (similar H_D1_/H_test_) and heteronymous Ia facilitation (similar H_Fac_/H_test,_ Figure 3) is based on the hypothesis that a constant conditioning stimulus would consistently excite GABAergic interneurons (D1 method) or motoneurons (HF), resulting in stable presynaptic inhibition of Ia afferents activity and additional stable heteronymous Ia facilitation, unless presynaptic inhibition of Ia afferents changes (Baudry & Enoka, 2009). Therefore, our results demonstrate similar presynaptic inhibition of Ia afferents and a concomitant stable heteronymous Ia facilitation during eccentric contractions at 20, 60 and 90°·s⁻¹. Heteronymous facilitation was assessed using short interstimulus interval scaling (0.5 ms) to reduce polysynaptic and postsynaptic inhibitory contamination and thereby enhance the selectivity of the measured facilitation (Meunier *et al*., 1994; Meunier & Pierrot-Deseilligny, 1998). This methodological approach is consistent with established practices in the literature and supports the robustness of our observations. Therefore, earlier predictions linking higher movement velocity to greater presynaptic modulation are not supported by our data, suggesting that other neural mechanisms are likely responsible for the observed reflex modulation. The observed reflex modulation may instead arise from postsynaptic mechanisms affecting motoneuron excitability.

### Recurrent inhibition on the velocity-dependent effect

To the best of our knowledge, the present study was the first to investigate the influence of angular velocity on recurrent inhibition. A key finding was that during eccentric contraction, increasing angular velocity was associated with a progressive enhancement of recurrent inhibition acting on SOL motoneurons (Figure 4). This observation is of particular interest because the pathway investigated does not represent recurrent feedback arising from the SOL motoneuron pool itself, but rather recurrent influences transmitted through an intermuscular spinal network involving quadriceps motoneurons (Katz & Pierrot-Deseilligny, 1999). Consequently, the present findings extend beyond the concept of local SOL self-regulation and suggests that recurrent interactions between motoneuron pools are sensitive to movement velocity.

The observation that increasing angular velocity selectively modulated a quadriceps-to-SOL recurrent pathway therefore suggests that adaptation to angular velocity is not restricted to mechanisms intrinsic to the SOL motoneuron pool. This interpretation is consistent with evidence showing that recurrent pathways are widely distributed across lower-limb motor nuclei and are strongly influenced by motor context (Katz & Pierrot-Deseilligny, 1999; Barbeau *et al*., 2000; Iles *et al*., 2000). In humans, heteronymous recurrent pathways from quadriceps to SOL are modulated according to postural and locomotor conditions and have been proposed to contribute to the coordination of locomotor transitions (Lamy *et al*., 2008). The present findings extend these observations by demonstrating that this pathway is also sensitive to angular velocity during eccentric contractions. From this perspective, the velocity-dependent increase in recurrent inhibition may reflect a functional reorganization of intermuscular spinal connectivity aimed at accommodating coordination demands as angular velocity increases.

This interpretation is further supported by the absence of velocity-related changes in SOL EMG activity. Because SOL EMG activity remained unchanged across angular velocities and EMG amplitude is commonly considered an indirect indicator of descending neural drive (Barrué-Belou *et al*., 2019), the increase in recurrent inhibition is unlikely to be solely explained by changes in descending neural drive. Instead, the dissociation between stable muscle activation and enhanced recurrent inhibition suggests a selective adaptation of inhibitory spinal interactions without substantial changes in overall motor output. This observation is consistent with the model proposed by Hultborn *et al*. (1979), whereby descending commands provide a relatively stable drive to α-motoneurons while recurrent circuitry contributes to task-dependent regulation of motor output under both spinal and supraspinal influences. Although the mechanisms underlying this modulation cannot be identified from the present data, the task-dependent regulation of recurrent pathways reported during locomotion suggests that supraspinal influences may contribute to the velocity-dependent reorganization of recurrent spinal connectivity observed here (Katz & Pierrot-Deseilligny, 1999; Iles *et al*., 2000).

The functional significance of this modulation may become particularly apparent at angular velocities approaching those encountered during locomotor activities. During walking, particularly during phases requiring eccentric control of the plantarflexors and during transitions from slow to faster gait, efficient movement execution depends on coordinated interactions among multiple lower-limb motoneuron pools. In this context, increased recurrent inhibition may represent an adaptive mechanism through which spinal networks adjust intermuscular connectivity as locomotor demands increase. In addition, Renshaw cell pathways have been proposed to regulate network gain and stabilize motor output by limiting excessive synchronization within motoneuronal populations (Windhorst, 1996; Uchiyama & Windhorst, 2007). Given that eccentric contractions are associated with greater motor unit synchronization than other contraction types (Duchateau & Enoka, 2016), the increase in recurrent inhibition observed during eccentric contractions at higher angular velocities may help prevent excessive synchronization within distributed motor networks, thereby contributing to stable motor output. Overall, the velocity-dependent increase in recurrent inhibition supports the view that recurrent spinal circuits are functionally organized according to task demands. Rather than reflecting a simple adjustment of local inhibitory feedback within the SOL motoneuron pool, the present findings suggest that increasing movement velocity is accompanied by a reconfiguration of inhibitory interactions acting on SOL motoneurons through distributed spinal networks. These findings highlight an important role of recurrent inhibition in the functional organization of intermuscular spinal connectivity and suggest that Renshaw circuitry contributes to the adaptive coordination of lower-limb motor pools during changes in angular velocity.

### Methodological considerations

To assess recurrent inhibition, we selected the heteronymous recurrent inhibition paradigm originally described by Meunier *et al*. (1994), as it represents the most appropriate approach for addressing our specific hypotheses. Given the distributed organization of recurrent projections across lower-limb, heteronymous measures may provide a more functionally relevant representation of recurrent spinal interactions. In the present study, this approach was especially appropriate because it allowed us to investigate whether increasing angular velocity during eccentric contraction modulates interactions between motoneuron pools (Barbeau *et al*., 2000) rather than the exclusive local regulation of SOL motoneuron excitability. Although a homonymous paradigm could theoretically provide a more direct assessment of recurrent inhibition acting on the SOL motoneuron pool, heteronymous approaches are widely employed in human neurophysiology research because of methodological limitations associated with homonymous paradigms. In particular, mixed nerve stimulation may introduce volitional-wave contamination, afterhyperpolarization effects and post-activation depression following the conditioning stimulus (Mazzocchio & Rossi, 1989; Rossi & Mazzocchio, 1991; Özyurt *et al*., 2019), all of which can influence the interpretation of conditioned reflex responses.

Other analytical methods including peristimulus time histograms and peristimulus frequencygrams allow a more precise estimation of the latency and duration of recurrent inhibition while reducing contamination from parallel reflex pathways (Brinkworth & Türker, 2003; Özyurt *et al*., 2019). The M-only method, which consists in isolating the direct motor response (M-wave) while minimizing or avoiding reflex (H-reflex) contributions, has also been proposed as a more selective approach, but it has been mainly applied during isometric contractions, where low-threshold motor units are preferentially and consistently recruited (Özyurt *et al*., 2019, 2020). In the present study however, measurements were obtained during eccentric contractions performed at different angular velocities, making the application of the M-only method impractical. In addition, the M-only approach requires invasive recordings, which were not compatible with our experimental design.

Antagonist activity, particularly through reciprocal inhibition mediated by Ia interneurons, may modulate the H-reflex pathway (Hultborn *et al*., 1976). It is therefore plausible that variations in antagonist activation during the experiment could influence agonist motoneuron excitability. However, the co-activation index was analysed to quantify the contribution of antagonist activation during plantarflexion tasks, and no main effect or interaction was observed across experimental conditions (all P values > 0.224). Taken together, these findings indicate that reciprocal inhibition arising from antagonist activity is unlikely to explain the observed variations in H-reflex responses.

Joint afferent input may contribute to increased Ib interneuron activity during eccentric contractions performed at high angular velocities (Pierrot-Deseilligny & Burke, 2005). However, during voluntary movement, this influence appears to be at least partly reduced by presynaptic inhibitory mechanisms acting at Ib afferent terminals, which are known to decrease the effectiveness of Ib inhibitory pathways (Lafleur *et al*., 1992). Therefore, while a contribution of this mechanism cannot be entirely excluded, it is unlikely to fully explain the reflex modulations observed in the present study.

Finally, although 90°·s⁻¹ cannot be considered an extreme angular velocity in experimental settings, the range of velocities tested in the present study (20, 60 and 90°·s⁻¹) covers low to moderate velocities that are representative of those encountered during functional movements. Notably, Mentiplay *et al*. (2018) showed that during ankle dorsiflexion in mid-stance, angular velocities increase with gait speed and can reach approximately 44–93°·s⁻¹ at walking speeds between ∼0.6 and 1.6 m/s (≈2.2–5.8 km·h⁻¹). This places our highest velocity condition within a physiologically relevant range. In this context, the increased recurrent inhibition observed at higher angular velocities may reflect neural control mechanisms engaged during natural locomotion. Importantly, the aim of this study was to investigate the modulation of Ia-to-motoneuron transmission behavior across a spectrum of angular velocities, rather than focusing exclusively on maximal contraction speeds, thereby capturing adaptations that are meaningful for everyday motor function.

## Conclusion

This study provides new insight into the influence of angular velocity on spinal inhibitory mechanisms during eccentric contractions. Our findings indicate that increasing angular velocity is associated with a reduction in Ia-to-motoneuron transmission, which appears to be primarily related to an increase in recurrent inhibitory activity. In contrast, presynaptic inhibition of Ia afferents remained unchanged across contraction velocities. Together, these results suggest a previously unrecognized role for recurrent inhibition in adjusting α-motoneuron discharge behavior in response to changes in angular velocity during eccentric contractions of SOL muscle. These findings advance our understanding of the neural mechanisms underlying the control of eccentric movements performed at different velocities, which are commonly encountered during functional human activities such as walking and running.

## ADDITIONAL INFORMATION

### Data availability statement

Data will be available online at the time of publication.

### Author contribution

JC: conceptualization, data acquisition, and writing original draft. KN: conceptualization and writing original draft. CL: data acquisition and writing original draft. JOL: data acquisition and writing original draft. TC: conceptualization and writing original draft. MJ: conceptualization, writing original draft.

